# Aged human iPSC-RPE organoid cultures display hallmarks of drusen formation

**DOI:** 10.1101/2021.10.12.463899

**Authors:** Lena Mesch, Natalia Pashkovskaia, Virginia Cora, Selin Pars, Serena Corti, Madalena Cipriano, Peter Loskill, Elod Koertvely, Stefan Kustermann, Marina Mesquida, Alexander Kleger, Stefan Liebau, Kevin Achberger

**Affiliations:** Institute of Neuroanatomy & Developmental Biology (INDB), Eberhard Karls University Tübingen, Österbergstrasse 3, Tübingen, Germany; Department of Biomedical Engineering, Sciences and Technologies, Eberhard Karls University Tübingen, Österbergstrasse 3, Tübingen, Germany; Roche Pharma Research and Early Development, Immunology, Infectious Diseases and Ophthalmology (I2O) Discovery and Translational Area, Roche Innovation Center Basel, F. Hoffmann-La Roche Ltd., Basel, Switzerland; Roche Pharma Research and Early Development, Pharmaceutical Sciences, Roche Innovation Center Basel, F. Hoffmann-La Roche Ltd, Basel, Switzerland; Department of Internal Medicine I, University Hospital Ulm, Ulm, Germany; Center for Neurosensory Systems (ZFN), Eberhard Karls University Tübingen, Tübingen, Germany

**Keywords:** AMD, hiPSC, RPE organoids, TIMP3

## Abstract

Age-related macular degeneration (AMD) is among the most common causes of irreversible vision loss. Disease progression is strongly associated with age-related pathological changes of retinal pigment epithelial (RPE) cells, such as accumulation of intracellular lipid-containing cell debris, extracellular lipid-rich deposits (drusen) and collagen-rich basal laminar deposits. Current AMD models provide a limited understanding of the complex pathomechanisms, revealing the lack of adequate physiological human AMD models. In this study, we developed an *in vitro* model applicable for the exploration of AMD pathomechanisms and risk factors for AMD progression and drusen formation. Advanced 3D culturing technologies allow long-term cultivation of hiPSC-derived RPE organoids (RPEorg) for up to 360 days, which is the time frame necessary for the development of an AMD-like phenotype. Aged RPEorg exhibit hallmarks of AMD and age-related alterations such as increased autofluorescence, accumulation of lipid droplets, calcification, and the formation of extracellular clusters of the drusen-associated proteins such as apolipoprotein E (APOE) and tissue inhibitor of metalloproteinases 3 (TIMP3). Electron microscopy further reveals drusen-like extracellular deposits mimicking the signs of late drusen formation and AMD progression. In summary, our results demonstrate that hiPSC-derived 3D RPEorg provide a promising model to study age-associated RPE pathology and drusen formation. We show here that RPEorg are applicable for disease modelling studies and early stages of drug development and provide the opportunity to uncover inter-individual genetic and epigenetic factors that alter the course of the disease.

## Introduction

Age-related macular degeneration (AMD) is one of the most common visual disorders in the senior population [1],[2]. Patients are initially affected by central vision impairment, followed by complete and irreversible blindness in later stages of the disease [3]–[5].

In AMD, histopathological changes mainly occur at the interface between the basal lamina of retinal pigment epithelial (RPE) cells and the inner collagenous layer of the Bruch’s membrane (BrM). Besides thickening and disorganisation of the BrM, an excessive accumulation of extracellular matrix (ECM) proteins and lipids can be observed below the RPE [6],[7]. Extracellular deposits, known as drusen, emerge between RPE and BrM and can be subdivided into hard drusen (circular deposits, up to 60 μm in diameter) and large pathological soft drusen (ranging from 30 μm to more than 1000 μm with an indistinct border) [7]–[9]. Hard drusen are considered a physiological sign of aging, but their enhanced occurrence is regarded as a risk factor for progression towards a sight-threatening condition [10]. Size and number of drusen depositions thereby positively correlate with the disease progression [11],[12]. In advanced stages of AMD, RPE cell atrophy and choroidal neovascularization may be observed [13]; events that finally lead to RPE cell loss and degeneration of adjacent photoreceptors (PRs) [7],[10].

The exact pathomechanism of drusen formation still remains to be clarified, although several studies propose a central role of RPE cells [14],[15]. *In vitro* mono-cultures of primary porcine RPE cells are even able to accumulate drusen-like material autonomously [16].

Adequate model systems are obligatory to gain a better understanding on the mechanisms of the drusen formation and AMD pathogenesis. Since animal models often fail to reproduce the hallmarks of AMD (especially due to the absence of anatomical characteristics such as the macula/fovea) and are accompanied by ethical concerns, robust and scalable *in vitro* models of drusen formation and AMD pathogenesis are urgently required.

There are several *in vitro* RPE models available. For instance, primary cells, e.g., human or porcine-derived, show strong similarities to human RPE cells *in vivo*, but have restricted availability (human explants) or pose interspecies differences, e.g., porcine cells. Most importantly, human RPE cells are examined post-mortem, representing mostly later stages of the disease, hampering studies of early phases of drusen manifestation. The emergence of induced pluripotent stem cells (iPSCs) [17] revealed new opportunities in the field of AMD modelling. Thereby, human iPSC of healthy and diseased donors can be differentiated into retinal tissues such as the neural retina in the form of retinal organoids (RO) [18],[19], as well as RPE [20]–[22]. Numerous studies have shown the capabilities of these model systems to recreate genetic diseases and ocular disease phenotypes [23]–[26].

Since AMD pathologies are strongly associated with aging, long-term cell cultivation is rather essential for exploring the effects of aging on RPE cell maturation and for recreation of the pathophysiologic events of early and late AMD stages. So far, RPE cells for AMD modelling have been mostly cultured two-dimensional on cell culture plates for several weeks or months. However, prolonged cultivation is associated with a tendency to detach from the culture vessel or transdifferentiate towards a mesenchymal phenotype [27]–[29].

Three-dimensional (3D) cultivation of RPE cells is a possibility to counteract the drawbacks of adherent cultivation. The generation of so-called spheroids from singularized RPE cell suspensions generates a functional phenotype [30],[31], however, an enzymatic dissociation process is required, that can negatively impact the matured, differentiated phenotype of RPE cells [28],[32]. Therefore, a second organoid-like approach for the generation of 3D RPE organoids (RPEorg) was implemented in this study, allowing for the self-assembly of RPEorg during differentiation of 3D retinal organoids (ROs) [21],[33]. Those spontaneously formed RPEorg reach high levels of maturation, as they can be cultured for a long time (>300 days) without intermediary dissociation steps.

In this study, we cultured RPE organoids for up to 360 days and demonstrate that these cells possess several hallmarks of drusen formation and aging, such as an increase of autofluorescence, elevated extracellular accumulation of the drusen-associated proteins APOE and TIMP3, as well as lipid droplet accumulation. Our results suggest that self-assembling RPEorg provide an easily accessible model to study the pathomechanisms of aging, drusen formation and risk factors for AMD development.

## Results

### hiPSC-derived RPE organoids display hallmarks of mature RPE cells

3D RPE organoids (RPEorg) employed in this study self-organized during retinal organoid (RO) differentiation (**Fig 1a**). Due to their strong pigmentation, RPEorg can be easily discriminated from other retinal and non-retinal tissues from day 40 of differentiation. They are typically attached to ROs, from which they can be mechanically detached by cutting before propagation (**Fig 1b**). When dissociated, RPEorg generate adherently cultured RPE cells (**Fig S1**). After 4 weeks of cultivation, RPE cells derived from early RPEorg displayed hexagonal RPE morphology and major RPE hallmarks (**Fig S1 and S2**).

**Figure 1.**
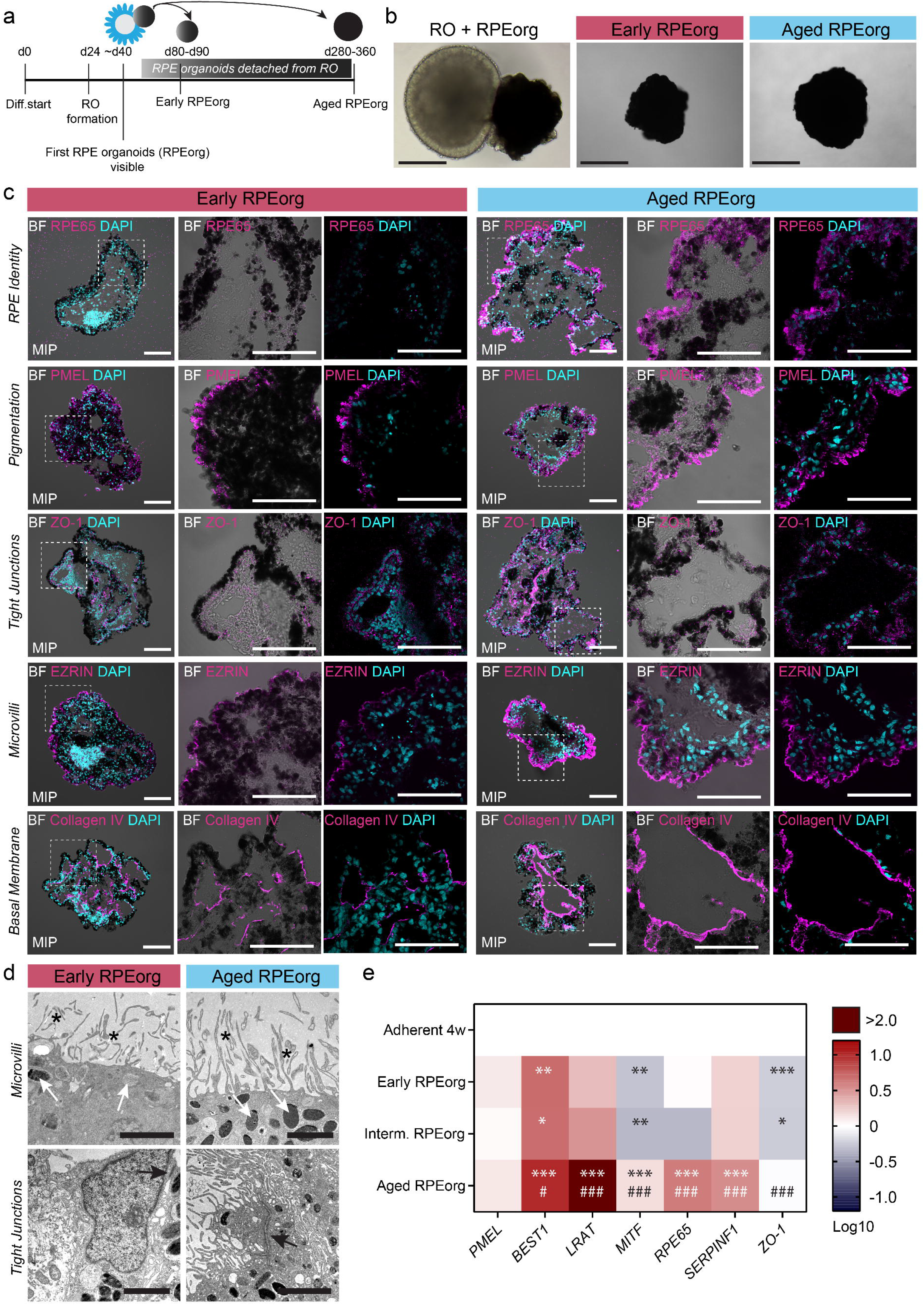
Aged RPE spheroids contain fully matured RPE cells. (a) Schematic depiction of RPE organoid differentiation and culture. (b) Brightfield images of RPE organoids attached to retinal organoids (left) and after dissection (middle, right). Scale bar: 1 mm. (c) RPE organoids (early and late stage) stained for RPE markers (magenta): (i) RPE65 (mature RPE, visual cycle), (ii) PMEL (pigmentation), (iii) ZO-1 (tight junctions), (iv) Ezrin (apical microvilli), (v) Col IV (basal membrane). White dotted squares in the first column per group indicate the magnified area depicted in the second and third image. DAPI = light blue. Scale bars: 100 μm. Maximum intensity projections are indicated as MIP. (d) TEM shows apical microvilli (*\**), melanosomes (*white arrow*) and tight junctions (*black arrow*) in early and late RPE organoids. Scale bars: 2,5 μm. (e) Heatmap revealing the gene expression of genes associated with RPE maturity, normalized to the housekeeping genes *GAPDH* and *RPS9* and shown relative to RPE cells dissociated from (early) RPE organoids and grown as a monolayer in a dish. The heatmap shows negative log-fold changes (blue) and positive log-fold changes (red). Expression of the following genes is presented: *PMEL* (pigmentation, melanosomes), *BEST1* (bestrophin-1), *LRAT* (Lecithin Retinol Acyltransferase, vitamin A metabolism visual cycle), *MITF* (Melanocyte Inducing Transcription Factor, early RPE marker), *RPE65*, *SERPINF1* (PEDF, secreted by RPE) and ZO-1. Statistical analysis represents the comparison between adherent RPE (*) and early RPEorg (#). n= 4 (Adherent RPE, early RPEorg, intermediate RPEorg) and n=12 (Aged RPEorg) per condition.

Next, early (day 80-90) and aged (day 280-360) RPEorg were characterized (**Fig 1c**). Immunohistochemical analysis of aged RPEorg displayed a strong expression of the RPE-specific 65 kDa protein (RPE65), an essential component of the visual cycle and consequently an important RPE identity marker, indicating an advanced maturation state. By contrast, RPE65 was barely detectable in the early stage RPEorg. Both early and late RPEorg were highly pigmented as demonstrated via brightfield imaging as well by staining for premelanosome protein (PMEL). Furthermore, RPEorg of both early and aged state, showed expression of the tight junction marker Zonula occludens-1 (ZO-1) at the cell junctions, the microvilli marker Ezrin, expressed on the outer surface of the organoid as wells as collagen IV as basal membrane marker. These findings were further substantiated by transmission electron microscopy. Here, melanosomes, tight junctions and apical microvilli could be identified in RPEorg of both analyzed maturation stages (**Fig 1d**).

In addition to the presence of RPE hallmarks in RPEorg, we also demonstrated a pronounced apical-basal cell polarization (**Fig 2**). The apical cell surface carried microvilli [34], while the basal surface of RPE cells produced collagen IV - a constituent of the basal membrane. The RPE basal membrane is one of the five layers of the Bruch’s membrane (BrM) and thus, not only characteristic for a functional RPE layer *in vivo*, but is also an essential component of an *in vitro* model of AMD, since disease-associated drusen are formed in the BrM.

**Figure 2.**
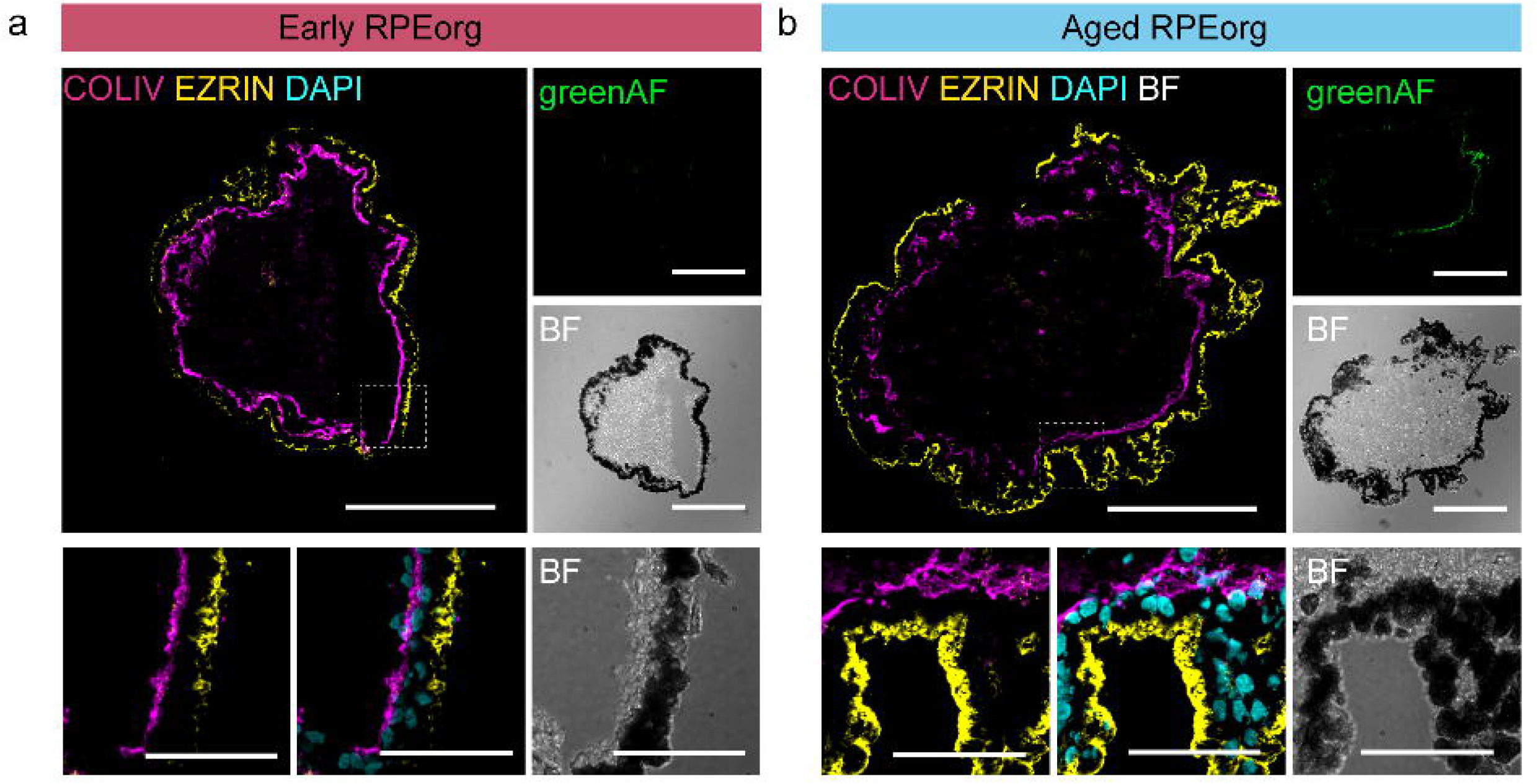
Early and aged RPE organoids show a prominent apico-basal polarity. (a) Early and (b) aged RPE organoids stained for collagen IV (COLIV, magenta, component of the basal membrane) and Ezrin (yellow, apical microvilli). White dotted squares in the upper image indicate the magnified area depicted in the images below. B = bright field. greenAF = green autofluorescence. DAPI = light blue. Scale bars: 200 μm (upper group) and 50 μm (magnified images below). All fluorescent images are maximum intensity projections (MIP).

One of the most important functions of RPE *in vivo* is to digest and recycle shed photoreceptor outer segments (POS). To investigate their phagocytotic potential, we fed FITC-labeled bovine rod POS to adherent RPE cells, early and aged RPEorg. After 8 h of treatment, both adherently cultured RPE cells (aRPE) and RPEorg displayed accumulations FITC/Rhodopsin double positive, suggesting that POS can be taken up by adherent and 3D RPE cells (**Fig S2a-b**, first panel). FITC+ accumulations were also positive for LAMP2 (**Fig S2a-b**, middle panel) and EEA1 (**Fig S2a-b**, last panel), indicating that aRPE and RPEorg are able to internalize POS respectively into lysosomes and early endosomes for further phagocytotic degradation (**Fig S2**).

Furthermore, the level of maturation of RPEorg and adherent RPE cultures were compared using qPCR. Gene expression of early (day 87-100), intermediate (day 192) and aged RPEorg (day 300-330) was compared to that of 4 weeks adherently cultured RPE cells (aRPE) (**Fig 1e**). We analyzed several essential RPE markers such as *PMEL* (premelanosome protein, marker for pigmentation), *BEST1* (bestrophin-1, RPE-specific calcium channel), *LRAT* (lecithin retinol acyltransferase, essential for the visual cycle), *MITF* (microphthalmia-associated transcription factor, RPE identity marker), *RPE65* (RPE-specific 65 kDa protein, visual cycle gene), and *SERPINF1* (serpin family F member 1, secreted by RPE cells). We observe higher expression of all these genes in aged RPEorg compared to aRPE. Early and intermediate RPEorg do not present apparent differences in marker expression, however, several markers (*PMEL*, *BEST1*, *LRAT*, *SERPINF1*) show higher levels in RPEorg of all ages than in adherent RPE (**Fig 1e**). Additionally, the increase of marker expression (except *PMEL)* between early/intermediate and aged RPEorg demonstrates a clear progress of maturation occurring between day 100/190 and day 300 (**Fig 1e**).

We confirmed RPE identity and maturity on morphological, protein and mRNA level in cultured hiPSC derived RPE organoids. Importantly, we also confirmed RPE-specific cell function by quantifying their phagocytotic potential and the expression of crucial visual cycle proteins.

### Drusen-associated proteins increase in aged RPE organoids

Sub-RPE deposits, reported to frequently produce focal aggregates known as drusen, are lipoproteinaceous waste products containing several extracellular matrix (ECM) components. Among them are apolipoprotein E (APOE) and tissue inhibitor of metalloproteinases 3 (TIMP3), which are present in drusen and upregulated in AMD [35]–[39].

To demonstrate the ability of RPEorg to model deposit formation and AMD-relevant protein accumulation, we assessed the expression of APOE and TIMP3 genes on protein and mRNA level. In comparison to early RPEorg, where TIMP3 was only weakly detectable, aged RPEorg showed a substantial higher TIMP3 expression (**Fig 3a, top panel**). In contrast, APOE expression remained unaffected and was present in early as well as in aged RPEorg (**Fig 3c**).

**Figure 3.**
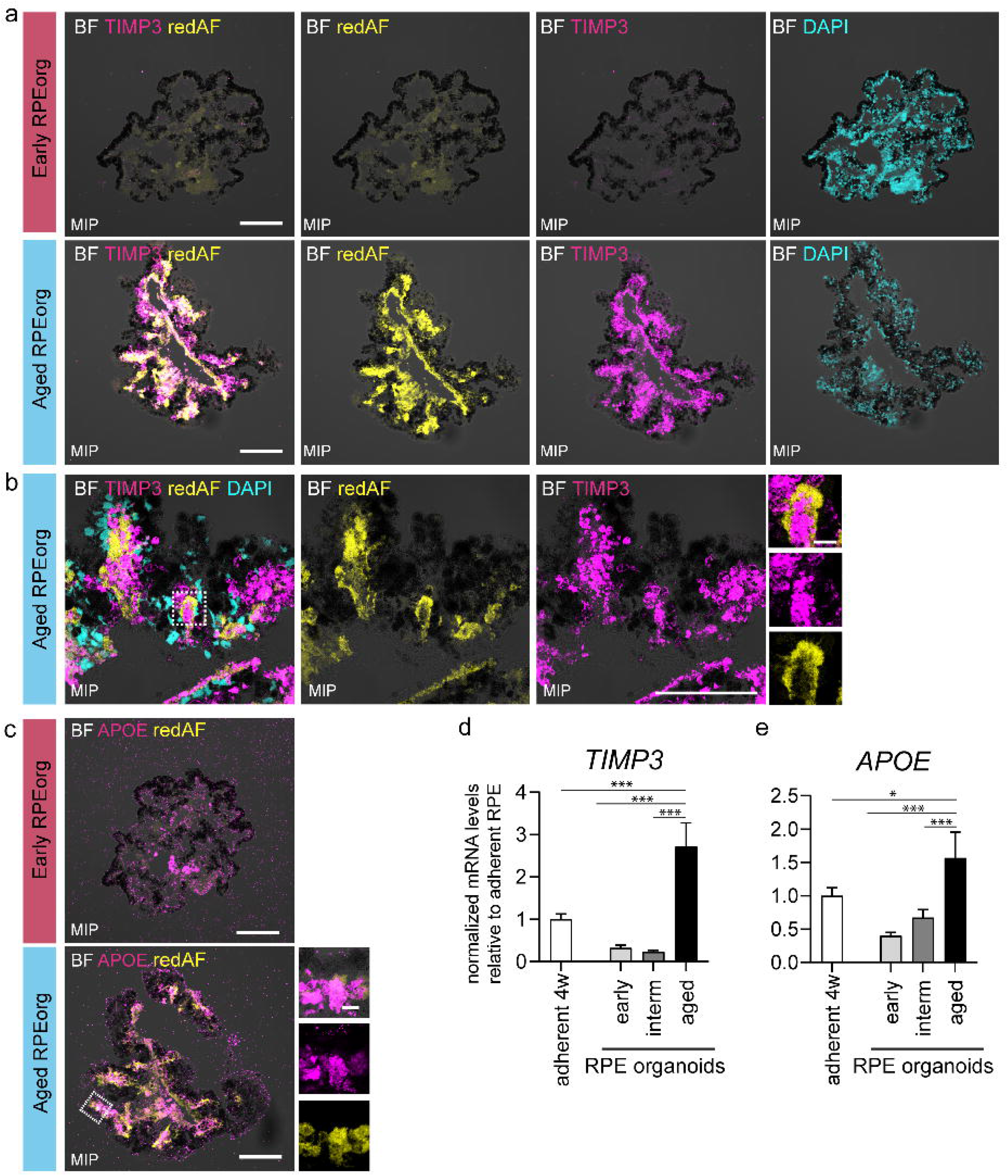
Drusen-associated markers and autofluorescence increase with RPE organoid maturation. (a-b) Cryosections of RPE organoids (early and late) were stained for drusen-associated proteins TIMP3 and imaged via confocal microscopy to assess red autofluorescence (redAF). (c) APOE staining and red autofluorescence (redAF). DAPI = light blue. Scale bars: large 100 μm. Magnified images (right) 10 μm. White dotted squares indicate the magnified area depicted in the images on the right. Images in a) and c) are maximum intensity projections (MIP). (d-e) mRNA expression of *TIMP3* and *APOE* normalized to *GAPDH* and *RPS9*. n= 4 (Adherent 4 weeks RPE, early RPEorg and intermediate RPEorg) and n=12 (Aged RPEorg) per condition. Mean + SEM.

TIMP3 spatial and temporal patterns correlated with red autofluorescence in early and aged RPEorg, although they largely did not overlap (**Fig 3a**). The presence of autofluorescence in RPE cells does not only occur *in vitro*, but is routinely used to detect the disease progression by fundus autofluorescence imaging in clinical examinations [40],[41].

The observed temporal expression of TIMP3 and APOE was confirmed by qPCR analyses (**Fig 3d**). Here, a robust increase in *TIMP3* expression was detected in aged RPEorg, compared to early and intermediate RPEorg and adherent RPE. *APOE* displayed a milder, yet significant, increase between aRPE and aged RPEorg, and a substantial increase from early to aged RPEorg.

Calcification in AMD can be traced back to the presence of crystallins (crystallin alpha a, CRYAA and crystallin alpha b, CRYAB) in sub-RPE deposits [7],[42],[43]. CRYAA and CRYAB are especially found in early/mid staged drusen [43] and could indeed be detected in both early as well as in aged RPEorg (**Fig S3a-b**). Although these proteins were detected in irregular patterns in our stainings of aged RPEorg, no clear increase or accumulation of any sub-RPE signal was observed. The investigation of RNA levels revealed a significant increase of *CRYAB* in RPEorg over time, paired with a decrease of *CRYAA* transcripts (**Fig S3d-e**). In comparison to adherent RPE culture, *CRYAA* mRNA levels were lower in RPEorg, whereas *CRYAB* levels were comparable.

Furthermore, ubiquitin (UBC) protein was highly expressed both in early and aged RPEorg (**Fig S3c**). This correlated with high mRNA levels throughout the RPEorg maturation, compared to UBC expression in aRPE (**Fig S3f**). Additionally, amyloid-β peptides, commonly associated with Alzheimer’s disease, have been previously reported to be implicated in drusen formation in AMD [44]–[46]. We found a strong and significant increase of amyloid-precursor-protein (APP) expression in aged RPEorg in comparison to early RPEorg and adherent RPE cultures (**Fig 3g**).

Other major components of AMD drusen are represented by extracellular matrix proteins, like Vitronectin (VTN), and complement factors, such as Complement C3, (C3), Complement C5 (C5) and Complement Factor H (CFH) [46]–[48]. In early and intermediate RPEorg, mRNA expression of *vitronectin* was poorly detectable (**Fig S3h**). In aged RPEorg however, the observed stronger vitronectin expression was comparable to that documented in adherent culture. Furthermore, mRNA of *C3*, *C5* and *CFH* could be detected in early as well as in aged RPEorg (**Fig S4a-c**). *C3* showed a significance increase between early and aged RPEorg, but the levels were overall lower than in adherently cultured RPE. *C5* levels did not change during RPEorg maturation but were overall higher than in adherent RPE. *CFH* was significantly higher in aged RPEorg than in early RPEorg, intermediate RPEorg or adherent RPE.

### RPEorg display neutral lipid accumulation and calcification

Having characterized the marker landscape in the RPE organoids, we sought to investigate the accumulation of drusen-characteristic lipids by performing whole-mount labelling with Nile Red, a fluorescent dye commonly used to label neutral lipid droplets [49]. To distinguish the Nile Red signal (red) from autofluorescence, green autofluorescence is displayed additionally (**Fig 4a**). Whereas early RPEorg displayed several areas of strong NileRed signal, aged RPEorg were characterized by an intense NileRed staining across their entire structure (**Fig 4a**). Nevertheless, quantification did not reveal a significant increase with aging (**Fig S5**).

**Figure 4.**
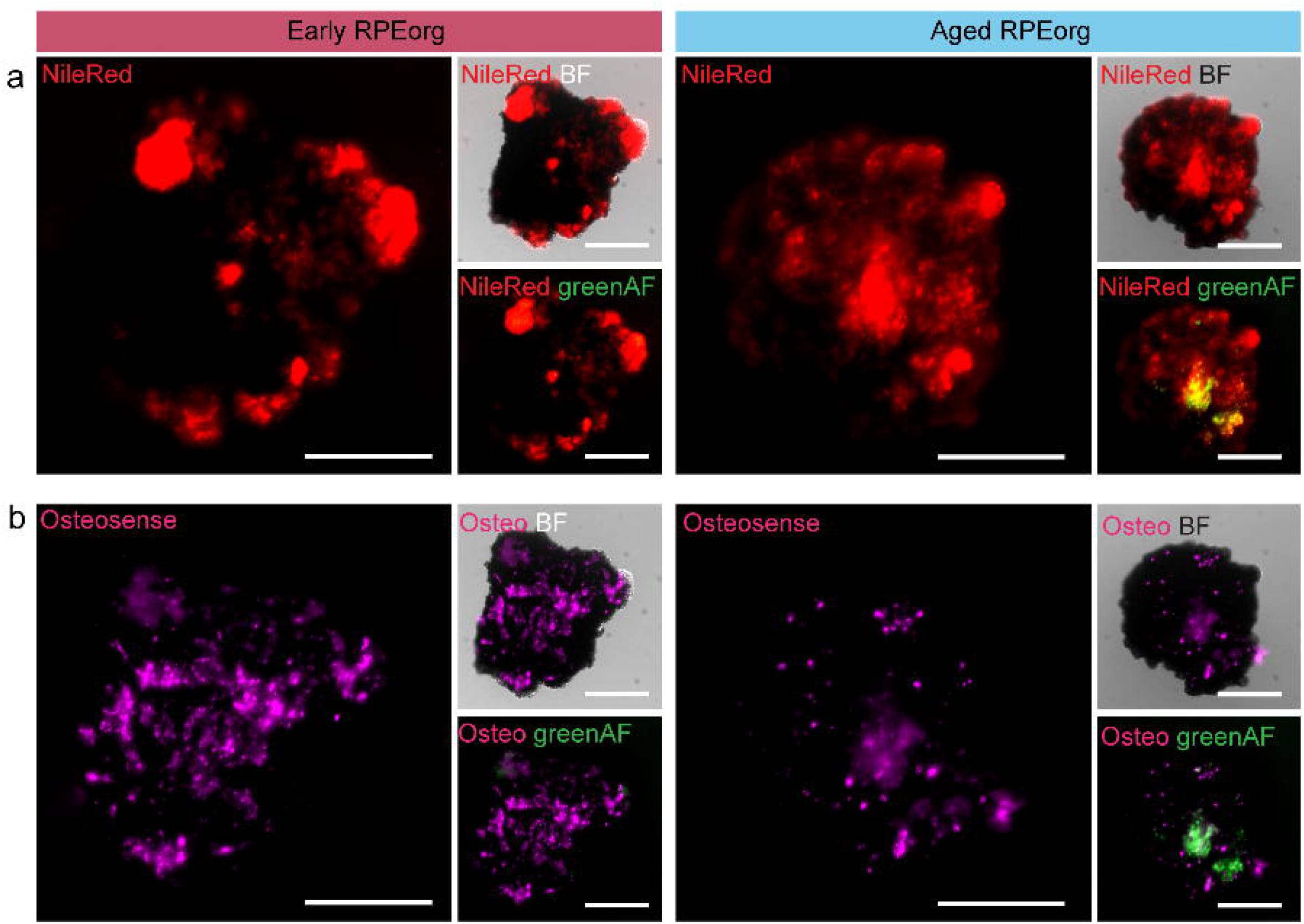
Matured RPE organoids contain strong patches of lipids and hydroxyapatite. Live cell RPE organoids labeled with (a) NileRed and (b) Osteosense. Green autofluorescence (greenAF). Scale bars: 100 μm

Calcification in form of hydroxyapatite (HAP) is another feature of drusen formation previously described for sub-RPE deposits [16],[50]. Detection of HAP in whole-mount stainings using the selective dye Osteosense revealed a strong signal not only in aged but also in early RPEorg (**Fig 4b**). Here, quantification surprisingly revealed an even higher level (**Fig S5**) confirming an early and ongoing calcification progress. Both presence of lipid droplets and calcification underline the presence of age and disease-relevant features in the RPEorg.

### Ultrastructural analyses reveal signs of basal deposits in aged RPE organoids

The accumulation of dark blue material below the outer cell layer of aged RPEorg in semi-thin sections implied the presence of lipophilic material as labeled by Richardson’s solution (**Fig 5a**). Electron microscopy analyses revealed various signs of deposits mainly in aged RPE organoids. These appeared as rather round structures enclosed by a membrane of either moderately electron dense interiors or smaller circular structures with a high electron-density (**Fig 5b, c**). In other regions, some clusters of wide-spaced collagen in combination with irregular structures of intermediate electron-density could be detected (**Fig 5d**). Additionally, scattered vesicular structures and basal infoldings were distributed within aged RPE organoids (**Fig 5e**). Similar structures have been characterized by TEM in literature [51]. In addition to homogenous electron-dense circular structures below the RPE, the accumulation of membranous debris and vesicular structures has been described [52],[53]. The presence of irregularly-shaped small structures with moderately dense lumina and basal deposits containing wide-spaced collagen were also suggested as potential characteristics of sub-RPE deposits [30],[47],[54]–[56].

**Figure 5.**
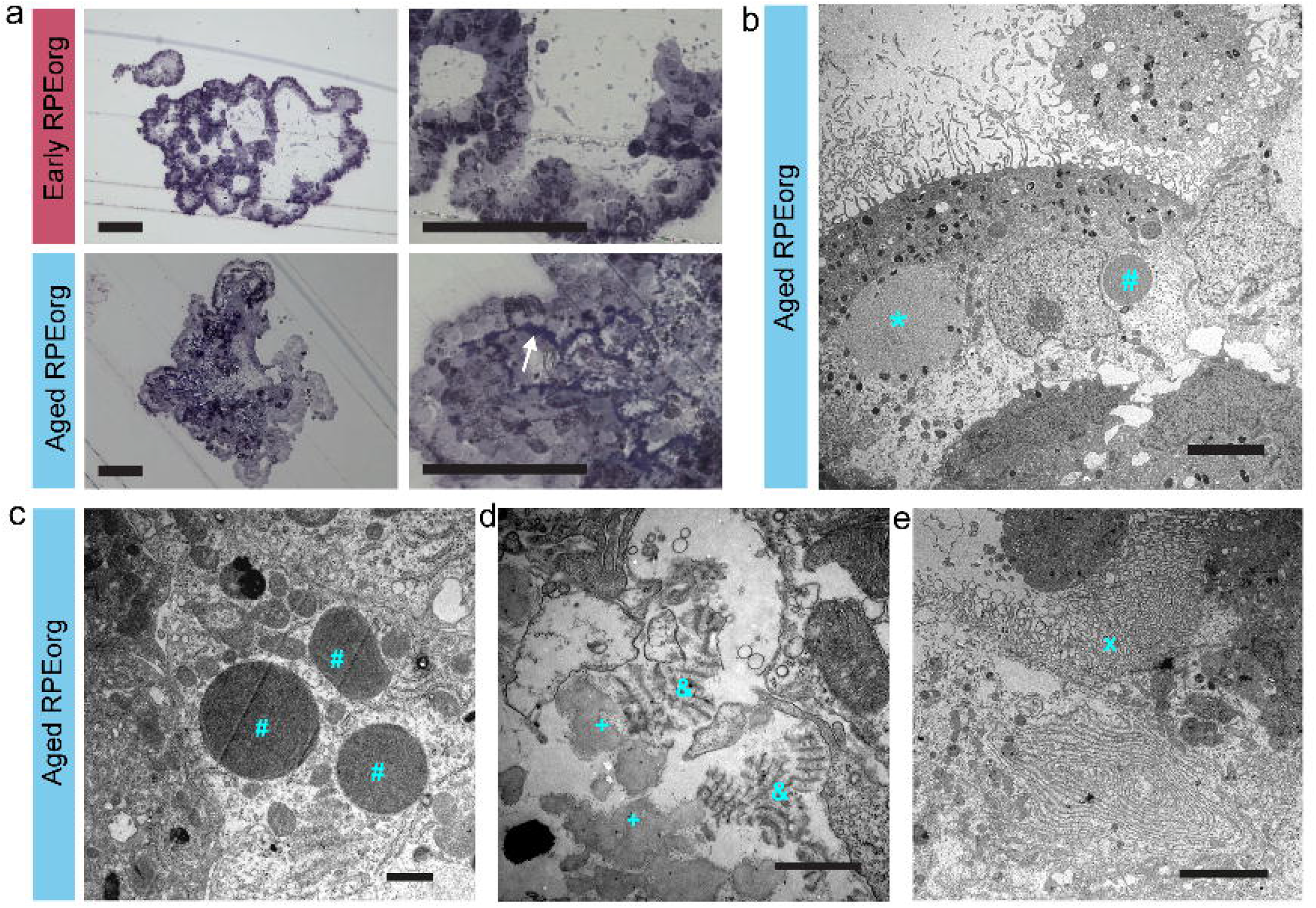
Ultrastructural analyses of RPE organoids. (a) Bright-field images of semi-thin sections reveal a strong accumulation of lipophilic material below the outer RPE cell layer in aged RPEs (white arrow). (b-e) Ultrathin-section transmission electron microscopy of aged RPEorg: Homogenous structures of intermediate electron density (*), circular, electron dense structures (#), wide-spaced collagen (&), small, irregular structures of intermediate electron-density (+) and vesicular structures (x). Scale bars: a) 100 μm, b, e) 5 μm; c d) 1 μm.

Taken together, our results indicate that aged RPEorg can recapitulate main AMD features, such as accumulation of the drusen-related proteins TIMP3, APOE, Vitronectin, C3 and CFH, and a gradual RPE cell autofluorescence increase. Moreover, the uniform expression overtime of CRYAA and CRYAB in RPEorg is comparable to the accumulation of these proteins in drusen during early and intermediate stages of AMD. Furthermore, aged RPEorg show several ultrastructural hallmarks of deposit and drusen formation underlying the relevance of this model of aged and AMD-related pathomechanistic research.

## Discussion

Over the last decade, 3D culture techniques have significantly improved the applicability of *in vitro* tissue models. Human stem cell-derived organoids and spheroids recapitulate the morphological and anatomical organization with high accuracy and can precisely model (patho-)physiological processes in human tissues and organs.

On the other side, classical 2D cultures of RPE cells often pose substantial disadvantages. Cultivation on artificial surfaces such as plastic or membranes renders an incomplete polarization and a poorly formed basal membrane. Most inconvenient is the fact that adherent cultures tend to detach or transdifferentiate after weeks or months of culture which makes them unsuitable for long-term investigations [16],[55]. In this work, we achieved to employ 3D cell culturing technologies to overcome the limitations of adherent RPE cultures creating a novel in vitro model for aging and AMD research.

RPE organoids, generated during retinal organoid differentiation from hiPSCs, allow a long-term cultivation of more than 300 days. They contain an organized layer of RPE cells, which display functional phagocytosis and form tight junctions. The presence of apical microvilli plus a collagen IV-positive membrane suggests a pronounced apico-basal polarization, an important prerequisite for RPE cell models [57],[58]. Moreover, the polarized RPE cell layer produces a basement membrane, resembling the human BrM. An important additional component of the BrM, the basal membrane of the choroidal vessels, could potentially be implemented by co-cultivation of RPEorg with endothelial cells or by assembloid fusion with vascular organoids [59]. Those approaches could presumably improve the formation of a more elaborated BrM and thus, facilitate studies on localization of sub-RPE deposits. All in all, our results suggest that RPEorg can mimic the architecture and function of the RPE layer *in vivo* and therefore provide a powerful tool for studying RPE functionality, as well as model diseases associated with RPE damage.

In this study, we showcased the applicability of the RPEorg model for AMD and especially the involvement of drusen in aging and disease. One of our major findings was the immense accumulation of TIMP3 in the central compartment of aged RPEorg. TIMP3 is a protein responsible for ECM remodeling by inhibiting matrix metalloproteases [60] and angiogenesis via VEGFR2 [61]. While TIMP3 is found in healthy hard drusen, various reports revealed its expression in the disease-associated soft drusen [36],[43],[62]. It has been hypothesized that TIMP3 can contribute to the thickening of the Bruch’s membrane, a major pathological feature of AMD [63]. In RPEorg, the accumulation of TIMP3 is found largely below the RPE and in the center of the organoid appearing thus as small aggregates which indicate the formation of depositions. These findings are supported by other drusen-associated markers such as APOE, CRYAA, CRYAB and Ubiquitin, which frequently form patches in the center of the organoid.

In clinical settings, fundus autofluorescence imaging is routinely used for determination of disease progression of AMD [40],[41]. A striking feature of aged RPEorg (200 or 330 days) was the appearance of autofluorescent material below the RPE layer in red and green spectrum. These autofluorescent granules did not directly colocalize with TIMP3-positive aggregates but were found in close proximity. This could correspond to intracellular lipofuscin granules, suggesting a sign of active RPE metabolism and aging [63],[64], but also to BrM and extracellular drusen [65],[66].

Amongst the other major components of drusen material are esterified or unesterified lipids [67]. In line with that, NileRed was employed to assess the occurrence of lipid accumulations in RPEorg. Our investigations revealed strong NileRed-positive structures in young, as well as in aged RPEorg, but did not delineate a significant effect during aging. A similar observation was made with respect to hydroxyapatite structures, labeled by Osteosense. Calcification has been described as an essential process during drusen biogenesis in AMD patients [47],[68]. Osteosense-positive patches were found in young as well as in aged RPEorg, indicating ongoing calcification during RPEorg cultivation. This is in accordance with previous *in vitro* studies showing early sings of calcification taking place in adherently cultured RPE [16].

Electron microscopy imaging showed spheroidal, electron dense (therefore presumably lipophilic) material inside and below RPE cells. Although this dense material strongly resembled hard drusen found in human eyes, it was with a diameter around 10 μm smaller than hard drusen *in vivo*, which can exhibit a size up to 63 μm [47],[63]. Additionally, accumulations of collagenous material and small amorphous structures were found in aged RPEorg and rather resembled soft drusen and basal linear deposits found in AMD patients. Furthermore, numerous membranous vesicles were observed in the center of aged RPEorg. *In vivo*, they are associated with the formation of soft drusen [47]. We conclude that aged RPEorg show several molecular and morphological features of age-related hard drusen formation, as well as early signs for the formation of soft, AMD-associated drusen. A potential explanation for the accumulation of disease-associated material in bona fide healthy cell lines might be related to the long culture time as well as the 3D structure of the RPEorg. While an organized (mono)layer of RPE cells surrounds the RPEorg, the interior is mostly cell free, containing ECM components or liquids. Due to the lack of the choroid-like blood stream system inside the organoids, waste products are potentially accumulating faster than in a healthy setting. In addition, the potentially hypoxic microenvironment in the center of the RPEorg might mimic the pathophysiological conditions also occurring in AMD, where the transport of oxygen is particularly hampered due to the drusen-caused thickening of the Bruch’s membrane [69],[70]. Phagocytosis of apoptotic material in the center of artificially created RPE spheroids has been observed in previous studies [30],[71].

RPEorgs presented in this study develop spontaneously during retinal organoid differentiation and are therefore less controllable than artificially formed spheroids. However, in comparison to previous described RPE spheroid models [30],[31], our results suggest that spontaneous formation of RPE is crucial for the advanced RPE features, the formation of a basal membrane and the occurrence of AMD-like features. Here, future in-depth analysis via single cell-based RNA sequencing could provide crucial hints on the precise cellular composition and maturation stage in the future. Furthermore, an extended functional characterization of the RPEorg model by, for instance, showcasing the barrier properties of RPE cells could increase the significance of the model for barrier and drug testing studies. Typically, this is demonstrated by measurements of the transepithelial electrical resistance (TEER) or diffusion of fluorescent dextran solutions [72]–[74].

In this study, we presented an extended characterization of the RPEorg features as well as their ability to recreate hallmarks of AMD disease. We anticipate that RPEorg which are derived from patients with genetic risk factors for AMD will even show a higher magnitude of drusen-like depositions and remodeling of ECM. Similarly, studies using long-term supplementation of AMD-associated stressors, such as smoke extract [75], photo-oxidized photoreceptor outer segments [76], A2E, blue light or the long-term addition of complement factors [77]–[79] could be implemented. We expect that future experiments employing such compounds could offer further insights into the biogenesis of drusen and their contribution to the AMD pathology.

In summary, the RPEorg model described in this study represents a powerful *in vitro* setting of mature RPE cells and thus allows studies of aging as well as early and intermediate biogenesis of drusen-like deposit formation occurring in AMD pathology. Moreover, we expect that the usage of this model could envision new venues for drug target identification as well as render new applications in drug efficacy studies.

## Acknowledgements

We would like to thank Anamaria Bernal-Vergara for culture and histological sample preparation. We further acknowledge Ulrich Mattheus, Institute for Clinical Anatomy and Cell Analysis, University of Tübingen for his assistance in sample preparation for electron microscopy and Prof. Dr. Ulrich Schraermeyer, Centre for Ophthalmology, University Hospital Tübingen for providing access to the electron microscope. This research was partly funded by the Fortüne programme (2586-0-0 to K.A.) of the University Tübingen, Germany and the DFG (DFG LI 2044/4-1, DFG LI 2044/5-1 to S.L.).

## Material & Methods

### iPSCs culture

HiPSC lines were generated from keratinocytes of healthy donors as previously described [80]. They were tested for stem cell markers and germ-layer differentiation potential. HiPSCs were grown on tissue culture plates (Becton Dickinson, USA) coated with hESC-qualified Matrigel (BD Biosciences, USA) in FTDA medium [81] and passaged every 5 to 7 days. Differentiated cells were removed regularly by scraping. Cells were grown at constant levels of 5 % O_2_ and 5 % CO_2_ at 37 °C. All experimental procedures were performed according to the Helsinki convention and approved by Ethical Committee of Eberhard Karls University Tübingen (Nr. 678/2017BO2). All donors gave their informed consent.

### hiPSC-derived RPE organoid differentiation and culture

HiPSC derived RPE organoids were differentiated along with retinal organoids, as previously published [82] based on a protocol by Zhong et al.[18] with some minor modifications. At day 24 of retinal organoid differentiation, the tissue containing neural retinal fields was sectioned into regular pieces, lifted manually, and grown in suspension in B27-based retinal differentiation medium (BRDM) containing DMEM/F12 (3:1), 2 % B27 supplement (w/o vitamin A, Thermo Fisher Scientific, USA), 1x minimum essential media non-essential amino acids (NEAA, Thermo Fisher Scientific, USA) and 1x antibiotics antimycotics (AA, Thermo Fisher Scientific, USA). From day 40 the medium was further supplemented with 10 % fetal bovine serum (FBS, Thermo Fisher Scientific, USA) and 100 μM taurine (Sigma-Aldrich, USA). Cells were cultured at 37 °C, 20 % O_2_ and 5 % CO_2_ in ultra-low attachment plates or petri dishes.

Non-retinal tissues were removed repeatedly from the suspension of retinal organoids and RPE organoids. RPE organoids became clearly visible between day 40 and 60 of differentiation and were typically attached to ROs or non-retinal tissue. For their purification and enrichment, RPE organoids attached to a different type of tissue were isolated with micro scissors and harvested accordingly. Pure RPE organoids were collected immediately.

### Cultivation of adherent hiPSC-derived RPE

For the generation of 2D adherent RPE, pure 3D organoids were cut in small pieces with microscissors and treated with Accumax (Sigma-Aldrich, USA) for 45 to 60 minutes at 37 °C. After straining the cell suspension with 70 μm and 40 μm cell strainers, cells were plated on cell culture plates (48-well or 24-well) coated with 0.01 % Poly-L-Ornithine (Sigma-Aldrich, USA) for 1 h at RT and 20 μg/mL Laminin (Roche, Switzerland) for 4 h at 37 °C. RPE cells were grown in BRDM supplemented with 20 ng/ml EGF, 20 μg/ml FGF2 (Cell Guidance Systems, United Kingdom), 2 μg/ml heparin (Sigma-Aldrich, USA), and 10 μM Y-27632 (ROCK-inhibitor, Ascent Scientific, USA) until they reached confluency. Additionally, 10 % FBS (Thermo Fisher Scientific, USA) was added for 24 h after plating. Adherent RPE cells were split every four to eight weeks with Accumax (Sigma-Aldrich, USA) and cultured in BRDM without FBS.

### Immunocytochemistry

For immunocytochemistry of cryosections, RPE organoids were fixed in 4 % paraformaldehyde (AppliChem GmbH, Darmstadt, Germany) for 20 min at room temperature (RT) and subsequently equilibrated to 30 % sucrose (Sigma-Aldrich, USA) overnight at 4 °C. After organoid embedding in cryomatrix (Tissue-Tek O.C.T. Compound, Sakura Finetek, USA), frozen samples were cut to 14 μm sections using a cryostat.

Cryosections were rehydrated for 30 min with phosphate-buffered saline (PBS, no calcium, no magnesium, Thermo Fisher Scientific, USA). Blocking and permeabilization were performed for 1 h at RT with 10 % normal donkey serum and 0.2 % Triton-X in PBS. The primary antibodies were diluted in blocking solution and incubated overnight at 4 °C. Secondary antibodies were diluted in 10 % NDS with 0.1 % Triton-X in PBS and incubated for 1 h at RT. Washing steps after primary and secondary antibody incubation were performed three times with PBS. Samples were finally stained with Hoechst 33342 (Thermo Fisher Scientific, USA) for 10 min at RT and mounted in ProLong Gold Antifade Reagent (Thermo Fisher Scientific, USA). Antibodies used for immunocytochemistry are listed in supplementary table S1.

### Phagocytosis Assay

Bovine photoreceptor outer segments (POS) were labeled with FITC using the Pierce FITC Antibody Labeling Kit (Thermo Fisher Scientific). Adherent RPE cells were treated with ~ 5 POS per cell and 1 x 10^6^ POS were applied per RPE organoid. After incubation with FITC-POS for 8 h at 37 °C, aRPE and RPEorg were washed five times with PBS −/− prior fixation, cryosectioning (only RPEorg) and immunostaining.

RPE organoids were used at day 92 (early RPEorg) and day 323 (aged RPEorg) with n=3. Adherent RPE cells were cultured for 14 days before performing the phagocytosis assay.

### Live cell labeling

#### NileRed

For live cell labeling of lipids with NileRed (Thermo Fisher Scientific, USA), RPE organoids were incubated in a non-treated cell culture plate for 25 min at 37 °C in BRDM containing 1 μM NileRed.

#### Osteosense

Live cell labeling of hydroxyapatite (HA) was performed with 1 μM OsteoSense 680EX Fluorescent Imaging Agent (Perkin Elmer) in BRDM for 20 minutes at 37 °C.

During the incubation period with both Osteosense and NileRed, RPE organoids were further labeled with Hoechst 33342 (Thermo Fisher Scientific, USA). Prior imaging, organoids were washed once with PBS.

### Fluorescence Microscopy

Images of stained RPE organoid cryosections were taken with a laser-scanning microscope (Zeiss LSM 710, Carl Zeiss, Germany). Labeled RPE organoids were analyzed using a conventional fluorescence microscope (Carl Zeiss, Germany), image stacks were recorded using an Imager M2 Apotome1 (Carl Zeiss, Germany).

In **Fig 2 and 4**, autofluorescence was recorded using a convential fluorescence Zeiss microscope using filter sets for red (Excitation 550/25, Emission 605/70) and green (Ex: 470/40, Em: 525/50) spectra. For **Fig 3 and S4**,autofluorescence was recorded using a confocal microscope with an argon 488 laser and a filter sets for green spectrum (Ex: 470/40, Em: 525/50).

### Transmission Electron Microscopy

Ultrathin sections for TEM were prepared as previously described [83], with some minor modifications. Briefly, RPE organoids were collected and fixed in Karnovsky buffer (2.5 % glutaraldehyde, 2 % PFA, 0.1 M sodium cacodylate buffer, pH 7.4) for 2 h at RT for TEM. After fixation, the samples were washed three times for 10 min in 0.1 M sodium cacodylate buffer (pH 7.4), postfixed for 1 h in 1 % OsO_4_ at RT and washed again three times in cacodylate buffer. Dehydration was performed in 50 % ethanol for 10 min and 70 % ethanol for 10 min, followed by treatment with uranyl acetate for 1 h at RT. Samples were washed with 70 % ethanol and 99 % ethanol over night at 4 °C. After three incubation periods for 10 min in acetone and another three repetitions of 10 min in propylene oxide, samples were infiltrated with increasing ratios of Epoxy embedding medium (Sigma Aldrich, Germany) in propylene oxide (25 %, 50 %, 75 %), each for 1 h. Finally, samples were infiltrated with 100 % Epoxy embedding medium for 2 h at RT in flat molds and cured for 12 hr at 60 °C. Semithin sections (1 μm) and ultrathin sections (50 nm) were cut on a Leica Ultramicrotome (Leica, Germany). Semithin sections were stained according to Richardson’s (Azure 0,5% Methylene blue 0,5%, Borax 1%). Ultrathin sections were collected on pioloform-coated copper grids. Sections were examined with a Zeiss EM 900 electron microscope.

### Gene expression analysis

Total RNA was extracted using the RNeasy^®^ Micro Kit (Qiagen, Germany), following the manufacturers protocol and stored at −80 °C. 20 ng of purified RNA was used for cDNA synthesis. The BioMark™ HD (Fluidigm, USA) was used for high-throughput gene expression analysis. Related Taqman™ probes were obtained from Thermo Fisher Scientific, USA, indicated in Table S1. Expression was normalized to a scaled mean of the housekeeping genes *GAPDH* (Glycerinaldehyd-3-phosphat-Dehydrogenase) and *RPS9* (Ribosomal Protein S9). The following formula was used:

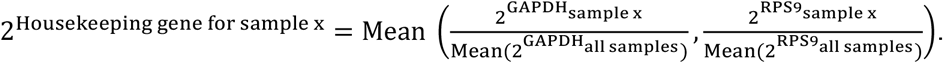

Graphs were set using GraphPad Prism (GraphPad Software, USA).

### Image Analysis

Fluorescence quantification was performed using ImageJ (imagej.nih.gov). Organoid area was manually selected, and intensity was quantified using the Measure plugin of ImageJ.

### Statistics

Statistical analysis was performed with GraphPad Prism 9.0 (Graphpad Software, USA) and statistical testing was performed using one-way ANOVA with Bonferroni post-hoc test. *p<0.05, **p<0.01, ***p<0.001.

## Author contribution

Conceptualization, K.A., P.L., S.L., S.K., A.K., E.K, M.M. M.C.; methodology, K.A., L.M., N.P., M.C, and E.K.; investigation, K.A., L.M., N.P., V.C., S.P., and S.C.; writing, K.A., L.M., N.P., E.K., A.K., M.M., M.C., S.L. and P.L.; resources, S.K.., S.L.., P.L. and E.K.; supervision: S.L., E.K. and K.A..

