## Supplementary Data for "Aged human iPSC-RPE organoid cultures display hallmarks of drusen formation"

### Supplementary figures

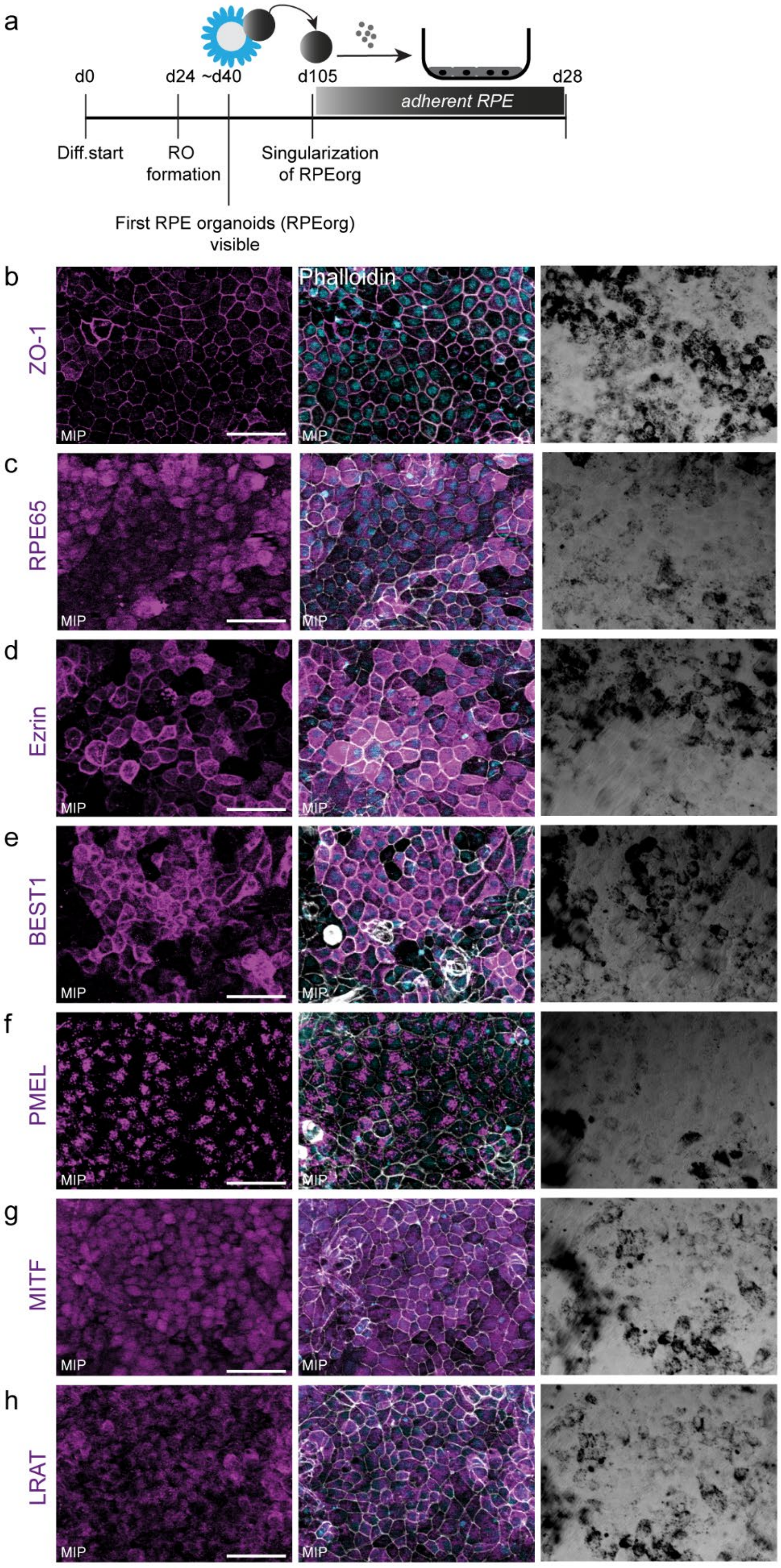


**Figure S1:** RPE grown as monolayer from RPE organoids cultured for 28 days.

(a) Generation of adherent RPE cultures from RPEorg. (b-h) Expression of RPE cell markers (magenta). Phalloidin = white. DAPI = light blue. Fluorescent images are maximum intensity projections (MIP). Scale bar 50 µm.

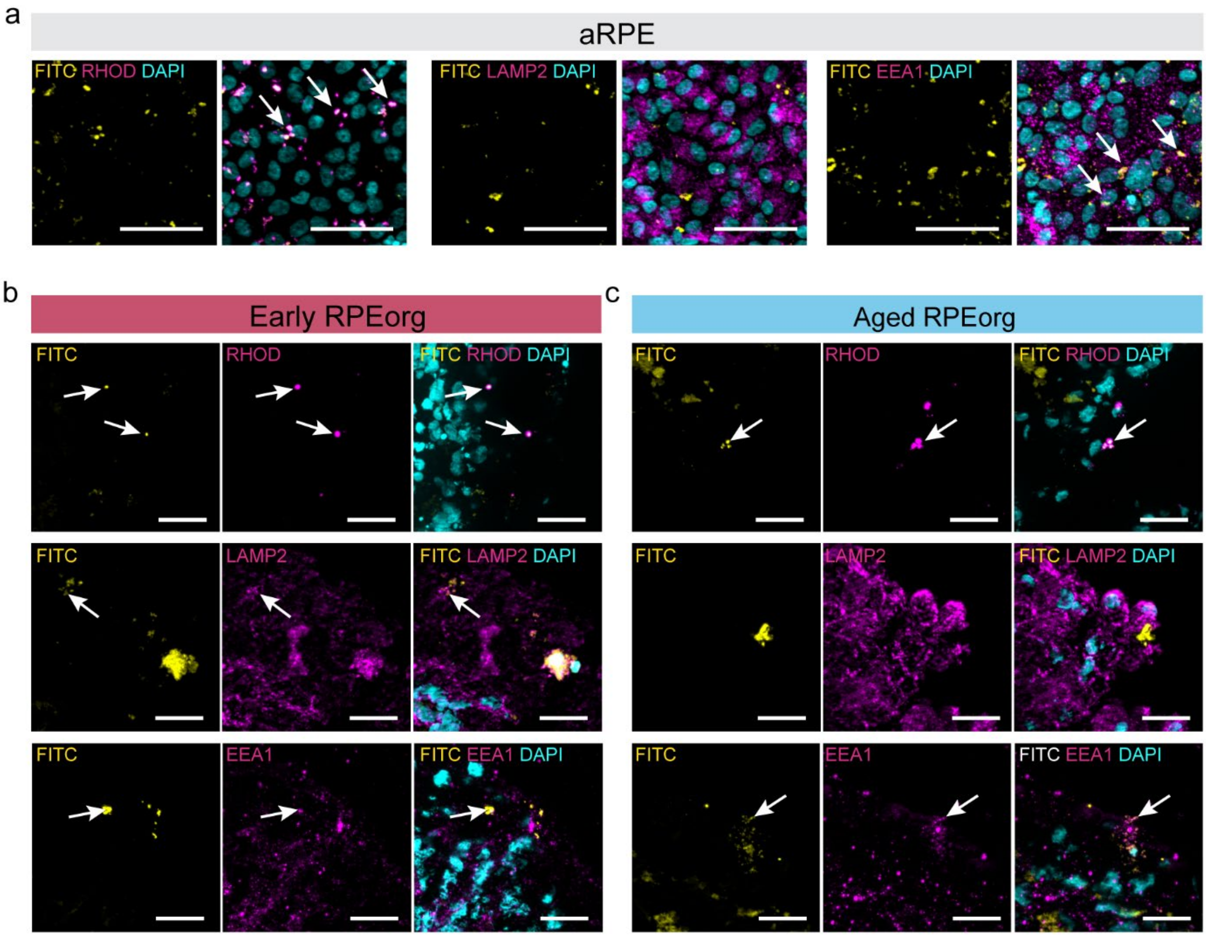


**Figure S2:** RPE organoids show phagocytotic activity.

(a) Adherent RPE and (b-c) RPEorg were treated with FITC-labeled bovine rod photoreceptor outer segments (POS, yellow) for 8 hours. Adherent RPE and cryosections of RPEorg were additionally stained for Rhodopsin (RHOD), LAMP2 and EEA1 (magenta). DAPI = light blue. All images are presented as maximum intensity projections (MIP). Scale bars: (a) 50 µm; (b-c) 20 µm.

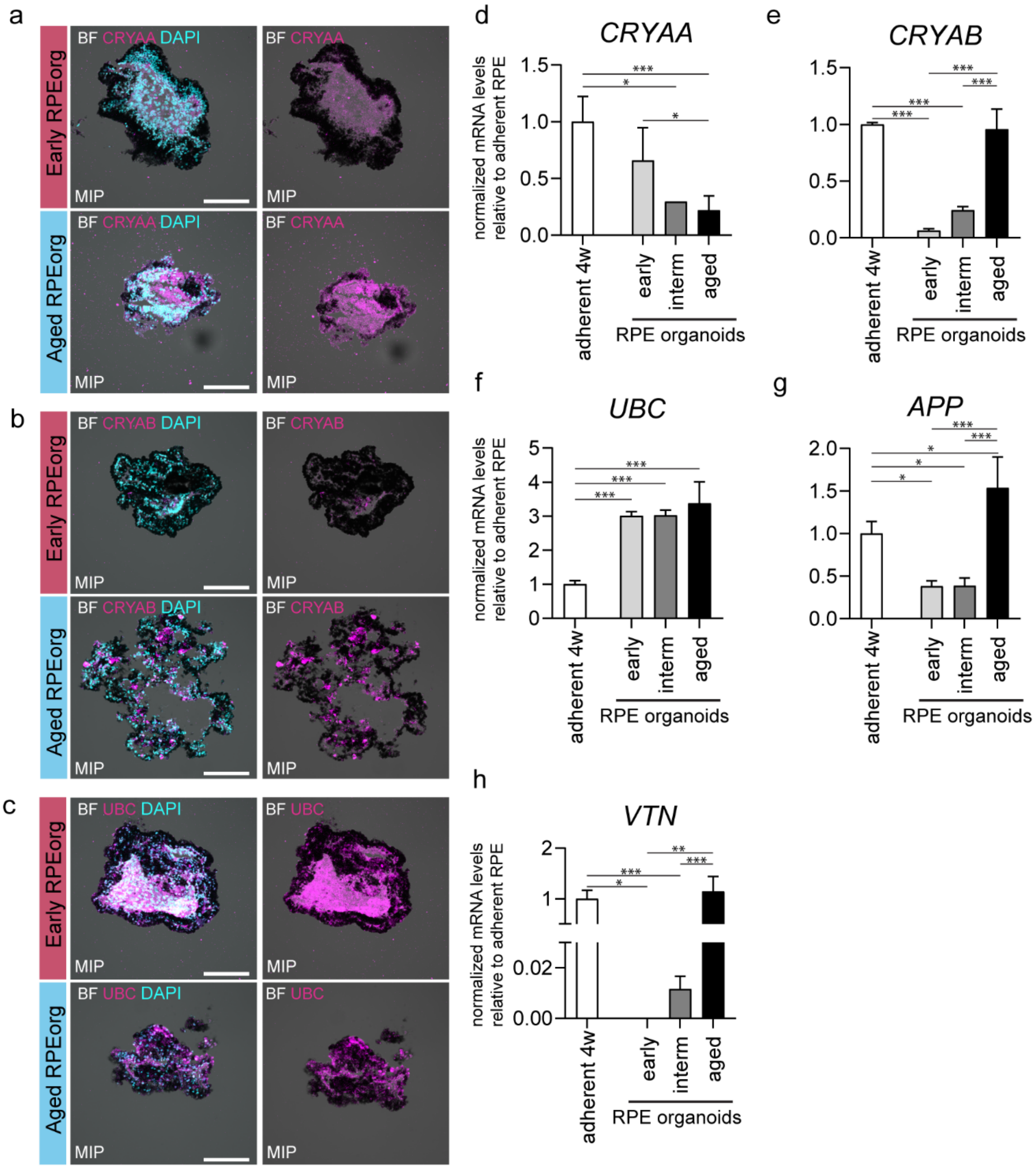


**Fig S3.** RPE organoids express drusen-associated proteins.

(a-c) Cryosections of RPE organoids (early and aged) were stained for drusen-associated proteins (a) α-crystallin (CRYAA), (b) β-crystallin (CRYAB) and (c) ubiquitin (UBC). DAPI = light blue. Scale bars: 100 µm. Images are maximum intensity projections (MIP). (d-h) RNA expression of *CRYAA, CRYAB, UBC, APP and VTN* normalized to *GAPDH* and *RPS9*. n= 4 (Adherent 4 weeks RPE, early RPEorg and intermediate RPEorg) and n=12 (Aged RPEorg) per condition. Mean + SEM.

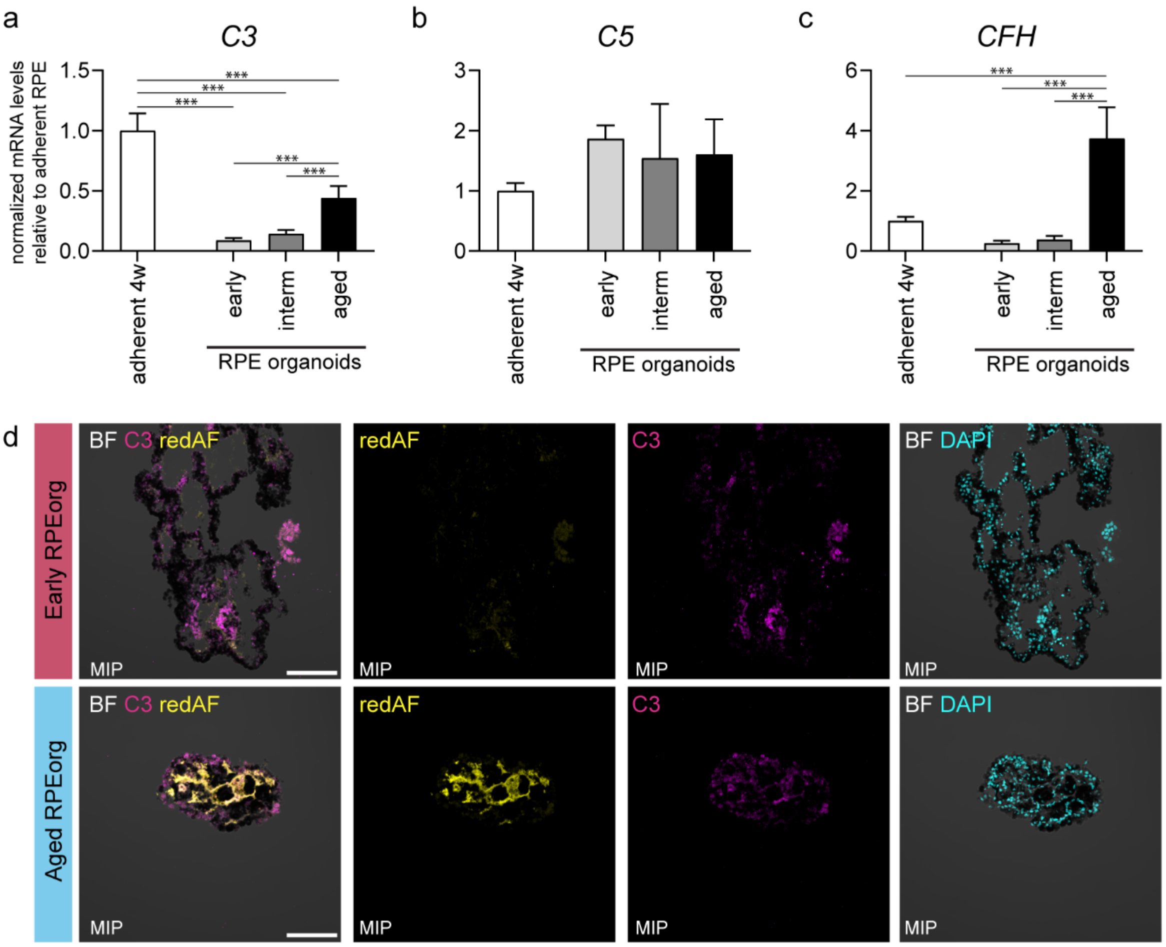


**Figure S4**. RPE organoids display presence of drusen-associated complement factors.

(a-c) RNA expression of *C3, C5* and *CFH* normalized to *GAPDH* and *RPS9*. n= 4 (Adherent 4 weeks RPE, early RPEorg and intermediate RPEorg) and n=12 (Aged RPEorg) per condition. Mean + SEM. (d) Immunostainings of early and aged RPE organoids with C3 (magenta) and red autofluorescence (redAF) were imaged via confocal microscopy. DAPI = light blue. Scale bars: 100 µm.

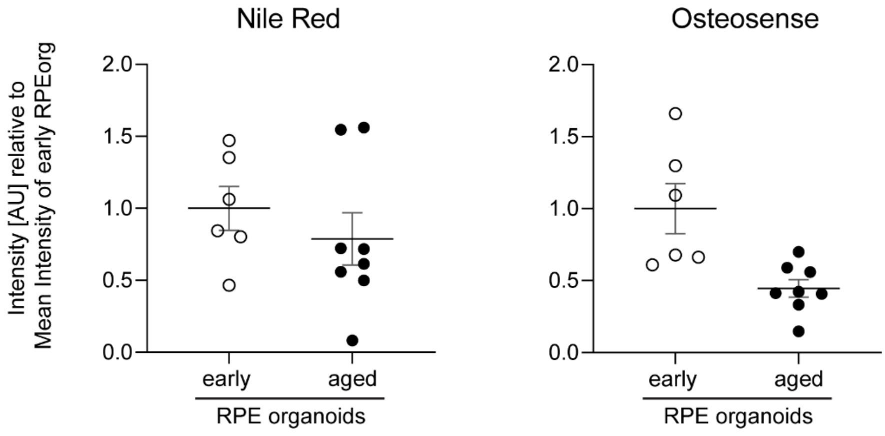

**Figure S5.** Quantification of neutral lipid accumulation (Nile Red) and calcification (Osteosense) in early and aged RPEorg.

Signal for Nile Red and Osteosense is displayed relative to the mean intensity of early RPEorg of each experiment. In total, n = 6 early RPE organoids and n = 8 aged RPE organoids were quantified.

**Supplementary table 1:** Antibodies for immunohistochemistry

| Primary Antibodies | Dilution | Ordering number | Company |
|---|---|---|---|
| ApoE | 1:150 | NB110-60531 | Novus, USA |
| C3 | 1:200 | PA5-21349 | Thermo Fisher Scientific, USA |
| C5b9 | 1:100 | NBP1-05120 | Novus, USA |
| Collagen IV | 1:50 | ab769 | Merck Millipore, Germany |
| CRYAA | 1:100 | CF505577 | OriGene, USA |
| CRYAB | 1:100 | CF500680 | OriGene, USA |
| EEA1 | 1:500 | 14-9114-82 | Thermo Fisher Scientific, USA |
| Ezrin | 1:200 | 3145S | Cell Signaling, USA |
| LAMP2 | 1:50 | sc18822 | Santa Cruz Biotechnology, USA |
| LRAT | 1:100 | LS-C416127-50 | LSBio, Seattle, WA, USA |
| Melanoma gp 100 | 1:200 | ab787 | Abcam, USA |
| RPE65 | 1:250 | ab78036 | Abcam, USA |
| TIMP3 | 1:200 | ab61316 | Abcam, USA |
| Ubiquitin | 1:100 | MAB1510-I-100UG | Merck Millipore, Germany |
| ZO-1 | 1:100 | 33-9100 | Thermo Fisher Scientific, USA |
| **Secondary Antibodies** | **Dilution** | **Ordering number** | **Company** |
| Donkey anti-Mouse Alexa Fluor® 488 | 1:500 | A-31571 | Thermo Fisher Scientific, USA |
| Donkey anti-Mouse Alexa Fluor® 568 | 1:500 | A-31571 | Thermo Fisher Scientific, USA |
| Donkey anti-Mouse Alexa Fluor® 647 | 1:500 | A-31571 | Thermo Fisher Scientific, USA |

**Supplementary Table 2** Taqman probes

| Gene name | Article number |
|---|---|
| *ApoE* | Hs00171168_m1 |
| *APP* | Hs00169098_m1 |
| *BEST1* | Hs04397293_m1 |
| *C3* | Hs00163811_m1 |

| | |
|---|---|
| *C5* | Hs01004342_m1 |
| *CFH* | Hs00962373_m1 |
| *COL4A1* | Hs00266237_m1 |
| *CRYAA* | Hs00166138_m1 |
| *CRYAB* | Hs00157107_m1 |
| *GAPDH* | Hs99999905_m1 |
| *LRAT* | Hs00428109_m1 |
| *MITF* | Hs01117294_m1 |
| *PMEL* | Hs00173854_m1 |
| *RPE65* | Hs01071462_m1 |
| *RPS9* | Hs02339424_g1 |
| *SERPINF1* | Hs01106937_m1 |
| *TIMP3* | Hs00165949_m1 |
| *TJP1* | Hs01551861_m1 |
| *UBC* | Hs05002522_g1 |
| *VTN* | Hs00940758_g1 |
